# *DMD* antisense oligonucleotide mediated exon skipping efficiency is affected by flanking intron retention time and target position within the exon

**DOI:** 10.1101/2022.10.28.514237

**Authors:** Remko Goossens, Nisha Verwey, Yavuz Ariyurek, Fred Schnell, Annemieke Aartsma-Rus

## Abstract

Mutations in the *DMD* gene are causative for Duchenne muscular dystrophy (DMD). Antisense oligonucleotide (AON) mediated exon skipping to restore disrupted dystrophin reading frame is a therapeutic approach that allows production of a shorter but functional protein. As DMD causing mutations can affect most of the 78 exons encoding dystrophin, a wide variety of AONs are needed to treat the patient population. Design of AONs is largely guided by trial-and-error, and it is yet unclear what defines the skippability of an exon.

Here, we use a library of phosphorodiamidate morpholino oligomer (PMOs) AONs of similar physical properties to test the skippability of a large number of *DMD* exons. The *DMD* transcript is non-sequentially spliced, meaning that certain introns are retained longer in the transcript than downstream introns. We tested whether the relative intron retention time has a significant effect on AON efficiency, and found that targeting an exon flanked at its 5’ by an intron that is retained in the transcript longer (‘slow’ intron) leads to overall higher exon skipping efficiency than when the 5’ intron is ‘fast’. Regardless of splicing speed of flanking introns, we find that positioning an AON closer to the 5’ of the target exon leads to higher exon skipping efficiency opposed to targeting an exons 3’-end.

The data enclosed herein can be of use to guide future target selection and preferential AON binding sites for both Duchenne and other disease amenable by exon skipping therapies.

## Introduction

Most gene transcripts undergo pre-mRNA splicing. In this process, the coding sequences of gene transcripts (exons) are joined together, while the non-coding intronic sequences are removed by the spliceosome in the form of lariat RNA. Introns are generally longer than exons and intron size ranges anywhere from less than 100 basepairs to over 100 kb. A notable example is the *DMD* gene, which encodes the dystrophin protein. The full-length transcripts are 2.21 Mb long, and introns make up 99.3% of the transcript size, varying in size between 107 bp and over 248 kb.

Mutations in the *DMD* gene that abolish production of functional dystrophin protein cause Duchenne muscular dystrophy (DMD), which manifests as progressive muscle wasting and loss of muscle function starting in young boys and leads to premature death, generally in the 3^rd^ or 4^th^ decade of life(Duan et al. 2021). The majority of mutations causing DMD are deletions of one or multiple exons, leading to a truncated DMD protein. A hotspot of reading frame disrupting *DMD* deletions is located between exon 43 and 55 (Aartsma-Rus et al. 2006; Tuffery-Giraud et al. 2009). Mutations in *DMD* can also cause Becker muscular dystrophy (BMD), which is characterized by a milder phenotype than DMD, but BMD mutations generally maintain the reading frame of *DMD(Aartsma-Rus et al. 2006)*. For about 55% of *DMD* mutations in DMD patients, it would be feasible to restore the reading frame by skipping a neighboring exon using antisense oligonucleotides (AONs)(Bladen et al. 2015). Currently, there are 4 FDA approved AON therapies for DMD, all of which target an exon in the exon 43-55 hotspot region(Schneider and Aartsma-Rus 2021). The purpose of these AONs is to induce exon skipping of exons 45, 51 or 53, which when applied to patients with a compatible lesion in *DMD*, can restore the reading from to generate a shorter, partially functional dystrophin protein as seen in BMD. However, with the currently approved AONs only about 30% of DMD patients can potentially be treated, meaning that a majority of DMD patients do not yet benefit from the current therapies. The development of different AONs necessary to skip the entire range of skippable *DMD* mutations is however a time consumable and difficult endeavor.

While AON design is guided by practical knowledge available in the scientific community(Aartsma-Rus et al. 2009), it is still unclear what truly designates a good targeting site for an AON to induce exon skipping. While studies have been published which attempt to catalogue the skippability of various *DMD* exons(Wilton et al. 2007), these results cannot be easily extrapolated as experimental design of the AONs, as well as coverage per exon, varies significantly. Furthermore, this data also showed that some exons are more difficult to skip than others, with the most difficult exons requiring combinations of AONS(Wilton et al. 2007). Based on retrospective analysis of effective and ineffective AONs, it is known that e.g. increased AON length, GC content and melt temperature play a role in AON efficacy(Aartsma-Rus 2012). Therefore it is difficult to compare which other parameters, such as targeting location, would make an AON reliably skip an exon.

Next to clinical necessity, using *DMD* as a model gene to study AON targeting is convenient as ample data is available about dystrophin transcript processing. These data indicate that *DMD* is non-sequentially spliced(Gazzoli et al. 2016), meaning that some introns are removed by the spliceosome earlier than an upstream intron which was transcribed before it by polymerase II (POLII). Based on our previously generated data, we classified that some of these introns are ‘slow’, while other introns are ‘fast’, concerning their splicing dynamics. This spatiotemporal model allows us to select sets of *DMD* exons flanked at either their 5’- and 3’-side by these ‘slow’ or ‘fast’ introns, and assess whether this characteristic influences the skippability of the exon. This model also allows us to assess whether targeting the AON itself more proximal or distal within the target exon is beneficial for exon skipping strategies, and whether this targeting location is dependent on the intron-class that flanks it on the target site.

We present here the first *in vitro* study where a large set of phosphorodiamidate morpholino oligomers (PMOs) with similar physical characteristics are used to determine optimal targeting of an exon to induce exon skipping. Our data indicates that exon skipping strategies have a significantly higher chance of successfully skipping an exon when the preceding 5’-intron is retained in the transcript for a longer amount of time. The speed at which the downstream 3’-intron is spliced does not seem to influence skipping efficiency. Furthermore, in the vast majority of targeted exons, there is a significantly higher efficiency of exon skipping when the AON is designed to target the proximal (5’) region of the exon. The data contained herein can be used to guide exonic target selection for Duchenne and other disease which are amenable for AON mediated therapies.

## Results

### Exon selection and PMO design

The non-sequential splicing of the *DMD* gene (Gazzoli et al. 2016), as well as the different splicing speed of its introns might have consequences for the efficiency of AON-mediated exon skipping strategies. Potentially, there might be a correlation between splicing dynamics and skippability of the exon. Previous work from our lab identified that various exons are flanked on their upstream 5’ splice site by an intron which is retained in the transcript relatively longer (slow intron) or shorter (fast intron), from here on, we will refer to these intron classes as 5’-Slow and 5’-Fast. Similarly, we denote the nature of the downstream 3’ intron as 3’-Slow and 3’-Fast to indicate that these introns were found to be retained relatively longer or shorter, respectively. For each of the potential combination of flanking intron classes (5’Slow + 3’Slow, 5’Slow + 3’Fast, 5’Fast + 3’Slow and 5’Fast + 3’Fast), we selected 3 out-of-frame exons of the *DMD* gene for which we designed PMO AONs (Figure 1A). To prevent efficiency bias from the physical properties of the AON, we designed panels of PMOs with similar physical properties, such as nucleotide count, GC content, melting temperature and potential free energy for forming secondary structures. We considered every possible 24/25-mer AON for eligible exons, resulting in 11.213 potential AONs. After calculating average GC content and melting temperatures, we only took PMOs, which were within 1 SD of the average GC content and melting temperature into consideration. After removal of AONs with unfavorable predicted free energy and secondary structure, we selected sets of PMOs for exons: 17, 21 and 70 (5’Slow + 3’Slow), 18, 22 and 50 (5’Slow + 3’Fast), 57, 65 and 67 (5’Fast + 3’Slow) and 51, 55 and 59 (5’Fast + 3’Fast) ((Figure 1A, Supplementary table 1), with larger exons being covered by a larger amount of PMOs (∼1 PMO/30 nucleotides). Overlap between PMO target sequences was prevented as much as possible.

**Figure 1:**
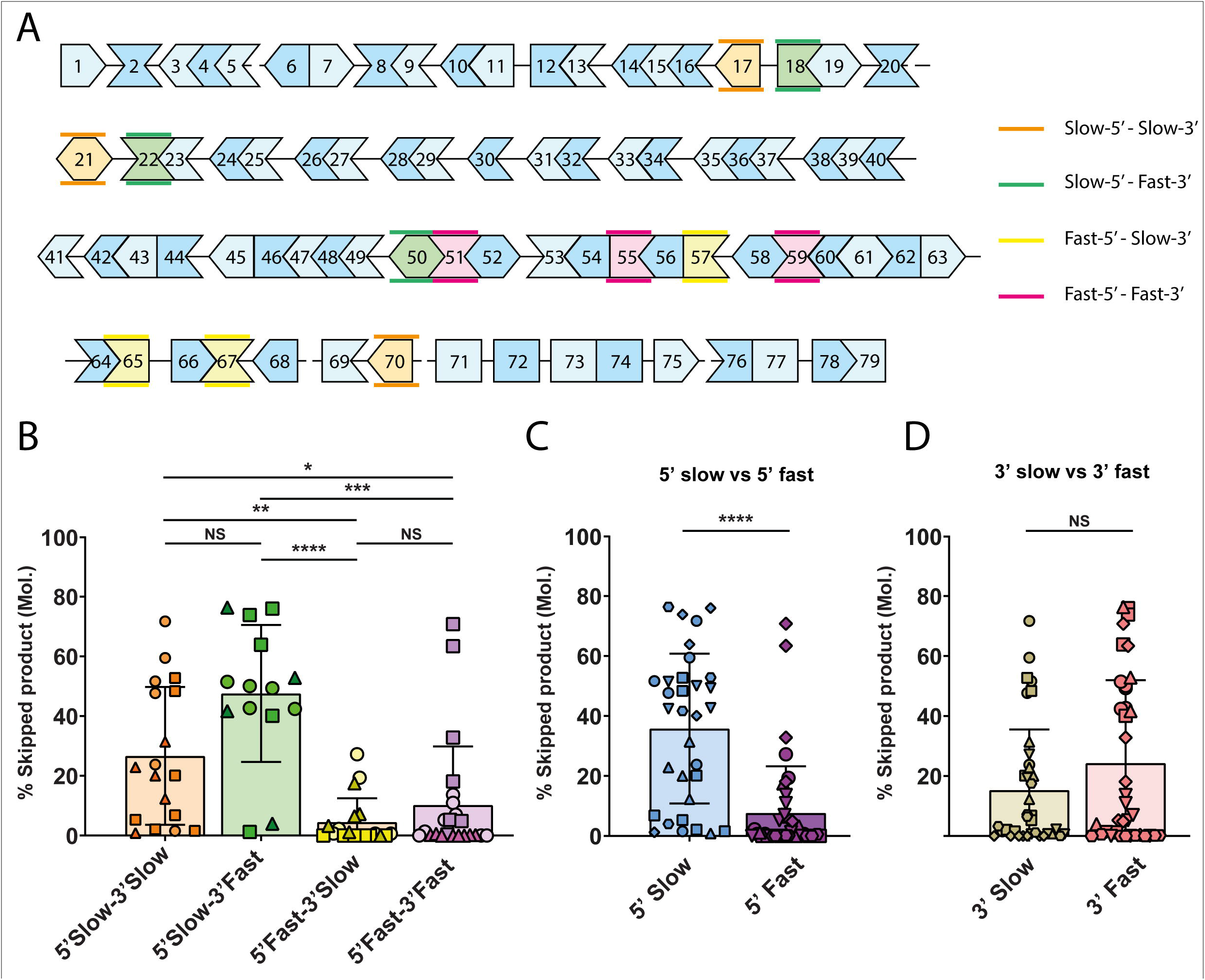
*DMD* exons flanked by a slow upstream intron are more susceptible to exon skipping. **A**: Graphical representation of the splicing order of introns in the *DMD* gene (adapted from Gazzoli et. al. 2016). Introns denoted by a line between exons are spliced slow, while exons shown directly adjacent are separated by ‘fast introns’. Different exon classes based on flanking introns are shown in orange, green, yellow and pink as indicated on the right. Different shades of blue are a visual aid with no relevance to the hypothesis. Reading frame continuity of the exon is indicated by the shape. **B**: Exon skipping efficiency of the *DMD* transcript with AONs for various exons as indicated in A. Each shape represents the average of two independent nucleofected samples, while each unique shape per bar corresponds to AONs targeting the same exon. Skipping efficiency was determined as the molar ratio of skipped product over total (full-length+skipped) product in an RT-PCR analysis using suitable primers for each exon. The exons belonging to the 5’Slow-3’Slow and 5’Slow-3’Fast classes show significantly higher exons skipping efficiency compared to 5’Fast-3’Slow and 5’Fast-3’Fast. **C**: Data presented in B was reanalyzed and grouped to separate exons based on their upstream intron class as indicated. 5’Slow exons show significantly higher skipping efficiency compared to 5’Fast. **D**: Data presented in B was reanalyzed and grouped to separate exons based on their downstream intron class as indicated. 3’Slow exons and 3’Fast exons show no difference in skipping efficiency. Error bars: SEM. (*: P value <0.05, **: P value <0.01, ***: P value <0.001, ****: P value <0.0001, NS: Not-Significant – One-Way ANOVA)

### Optimization of PMO delivery to myotube cultures

To facilitate reliable and reproduceable delivery of the PMOs to the control myoblast line we used electroporation in an Amaxa 4D nucleofector X unit with 16-well nucleocuvette strips (Aung-Htut et al. 2019). We validated the delivery of PMOs using this system in our cell line with a single exon 51 targeting PMO, using various buffer systems and pulse programs. We selected the condition with the highest detectable skipping of exon 51 using RT-qPCR (Supp. Figure 1), and used this condition (Buffer set P1, pulse program CM-137) for the remainder of the project. Electroporation of myoblasts still allows for differentiation to *DMD* expressing myotubes, as determined by RT-qPCR for myogenic markers *MYOG* and *MYH3* (Supp. Figure 1A), The foremost used method for semi-quantification of exon skipping PCR products is the Agilent Bioanalyzer 2100 using a DNA-1000 chip. To facilitate the high amount of generated samples, we compared the Bioanalyzer 2100 data with data generated by the Agilent Femto pulse system, which allows higher automated throughput of samples. Running the same RT-PCR sample on both systems showed no discernable differences in estimated exon skipping efficiency (Supp. Figure 1B).

### Exons with slow 5’-introns appear to be more skippable

We electroporated the set of PMOs targeting the 12 selected exons outlined above in HC myoblasts and allowed cells to differentiate to myotubes. After RT-PCR and analysis of exon skipping efficiency (Supp. Figure 2), we observed that there was a clear, statistically significant, difference in skipping efficiency between the different exon classes (Figure 1B). The 5’Slow-’3Slow and 5’Slow-3’Fast showed higher average skipping efficiencies than the 5’Fast-3’Slow and 5’Fast-3’Fast exon classes. No significant difference was observed between 5’Slow-3’Slow and 5’Slow-3’Fast, or between 5’Fast-3’Slow and 5’Fast-3’Fast. Confirming this, when reanalyzing the data and grouping of the exons solely based on splicing speed of their 5’-or 3’-flanking intron, we only observed a significant difference in skipping efficiency when grouping for the splicing speed of the 5’-intron (Figure 1C), but not the 3’-intron (Figure 1D). Together, this data shows that a longer retention of the upstream intron has a positive effect on exon skipping efficiency.

### An AON targeting the 5’ region within the exon is more efficient

To test whether the optimal target within an exon was determined by the presence (5’ or 3’) of slow introns, we designed another set of PMOs, now only using 5’Slow-3’Fast (exon 10, 14, 18, 22, 42 and 53) and 5’Fast-3’Slow (exon 9, 27, 52, 57, 65 and 67) exon classes (Figure 2A, Supplementary table 1). We aimed for 8 AONs per exon, positioning 4 AONs in the proximal 30% and 4 AONs in the distal 30% of the exon. Using these requirements, it was not feasible to select only out-of-frame exons while still adhering to our design criteria for PMO similarity. Therefore we selected 6 *DMD* exons for each class from both out-of-frame (exon 18, 22, 52, 53, 57, 65 and 67) and in-frame exons (exon 9, 10, 14, 27 and 42).

After delivery of this set of PMOs to HC myoblasts and differentiation to *DMD* expressing myotubes, we analyzed skipping efficiency of each PMO as before, using suitable RT-PCR primer sets for each targeted exon. The relative binding position of each of the AONs was calculated as the most proximal or distal exonic position possible for a 25-mer AON on an arbitrary 1-100 scale, and skipping efficiency was plotted on this coordinate for the corresponding PMO. When analyzing the skipping efficiency of PMOs in the 5’Slow-3’Fast category (Figure 2B) or 5’Fast-3’Slow category (Figure 2C) we observed a trend of higher skipping efficiency closer to the 5’ of the exon for almost all exons tested. This was confirmed by regression analysis of the skipping efficiency of each exon, which showed a negative slope for all skippable exons (Figure 2B, 2C and Supp. Figure 3). Exons not abiding this observation were exon 65 and 10, which showed no skipping for any of the individual PMOs tested (Supp. Figures 3 and 4).

**Figure 2:**
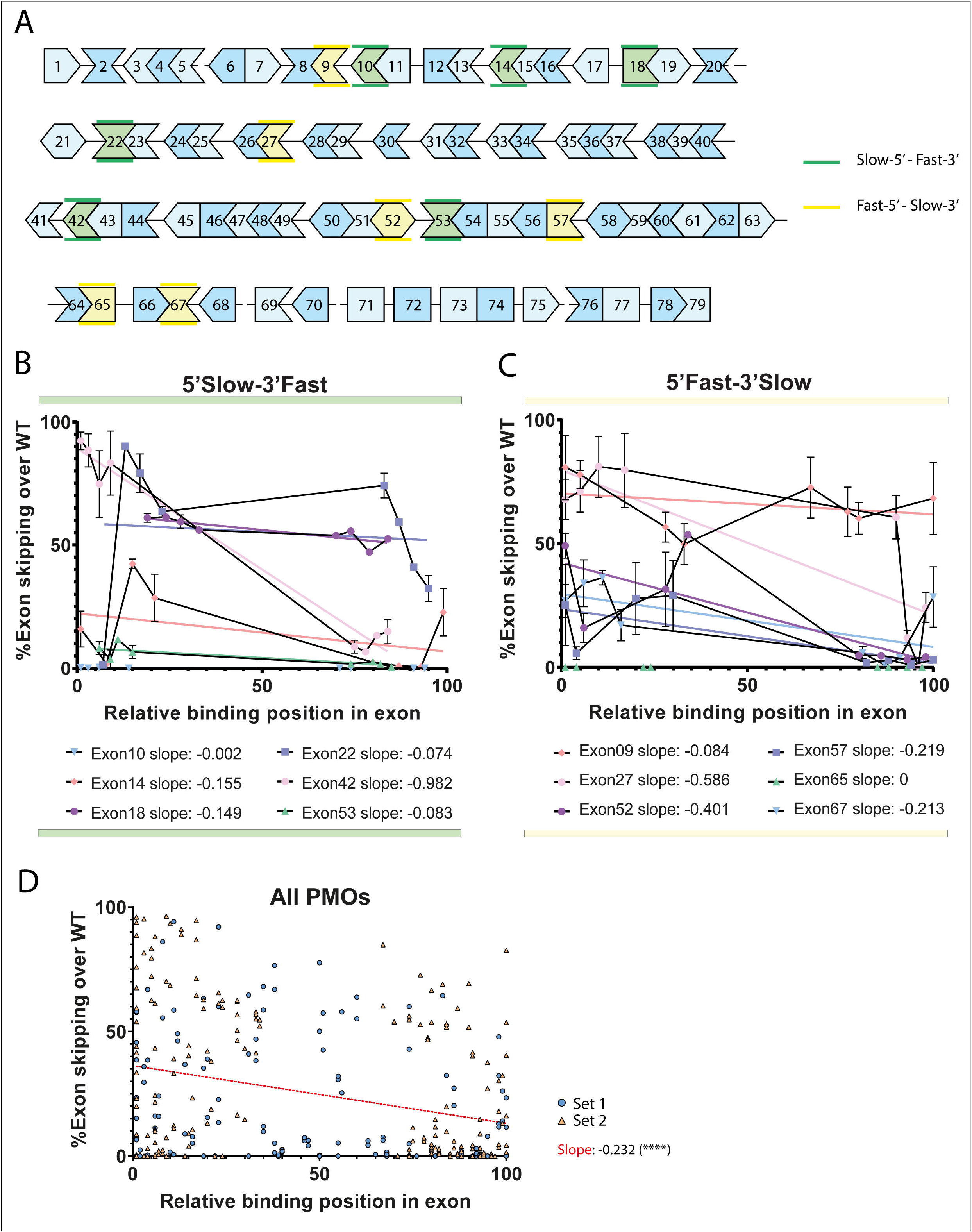
Targeting the 5’ of an exon leads to higher exon skipping efficiency. **A**: Graphical representation of the splicing order of introns in the *DMD* gene (adapted from Gazzoli et. al. 2016). Introns denoted by a line between exons are spliced slow, while exons shown directly adjacent are separated by ‘fast introns’. Different exon classes targeted for AON mediated exon skipping based on flanking introns are shown in green and yellow as indicated on the right. Different shades of blue are a visual aid with no relevance to the hypothesis. Reading frame continuity of the exon is indicated by the shape. **B**: Exon skipping efficiency of the *DMD* transcript with AONs for various exons from the 5’Slow-3’Fast flanking intron class as indicated in A. Individual points indicate the average of two independently nucleofected samples. The X-axis represents the possible window of targeting a 25-mer AON within the exonic sequence of the indicated exon, scaled from 0 to 100 for each exon to normalize exon size. Lines matching the symbol colors represent the results of a linear regression analysis of the skipping efficiency as a function of the targeting position. Individual slopes of each regression analysis are indicated below the plot. **C**: Exon skipping efficiency of the *DMD* transcript with AONs for various exons from the 5’Fast-3’Slow flanking intron class as indicated in A. Plots, axis and analysis description are identical to the plot presented in B. **D**: Reanalysis and summarizing plot of the *DMD* exon skipping efficiencies of all AONs used in figures 1B, 2B and 2C, as a function of their position within the exon as indicated on the X-axis. Circles indicate data from figure 1, triangles represent data from figure 2. Linear regression analysis (red line) shows a negative slope, indicating that in general, an AON targeting closer to the 5’-end of the exon will be more efficient at skipping the target exon than a more distally targeted AON. Error bars: SEM. (****: P value <0.0001 – Linear regression)

This correlation between AON targeting site and skipping efficiency was also observed when we combined the data of all PMOs used in this study (Figure 1 and Figure 2) and plotted the regression analysis (Figure 2D). We noted a significant negative trend towards the 3’ of the exon, indicating that in general, an AON targeting the 5’ of an exon has higher potential of being effective, regardless of the flanking introns splicing dynamics.

Out-of-frame transcripts might be unstable, causing them to be degraded by non-sense mediated decay (NMD), a cellular mechanism used to degrade mRNA molecules which contain premature stop-codons. Degradation by NMD could cause a bias when targeting both in-frame and out-of-frame exons. Indeed, there seems to be a trend to higher skipping efficiency for AONs targeting in-frame exons (Supp. Figure 3D). It should be stressed, however, that the sample size of out-of-frame exons is much larger in our analysis.

### Exon skipping can lead to transcript loss in control cells

We then aimed to investigate whether the low levels of exon skipping observed in control myotubes for exon 51 and 53 could be partially explained by the fact that these experiments would generate an out-of-frame *DMD* transcript, which could be a target for NMD. We electroporated sets of exon 51 or exon 53 targeting PMOs in two DMD patient derived cell lines. These cells harbor a deletion of *DMD* exon 48-50 (line 8036)(Mamchaoui et al. 2011) or exon 45-52 (line 6311)(Echigoya et al. 2019), making them amenable for reading-frame restoration by skipping of exon 51 or exon 53, respectively. When comparing the exon skipping levels in controls observed for exon 51 (Figure 3A) or exon 53 (Figure 3B) with the respective DMD cell lines (Figure 3C and 3D), it is apparent that the majority of targeting AONs result in higher exon skipping levels in DMD patient derived cell lines. This indicates that the detectable levels of exon skipping in control cells might be underestimated due to degradation of the skipped transcript. However, trends in AON efficiency remain the same between control and patient cells indicating that the relative skipping efficiency can still be deduced by skipping out-of-frame exons in control cells. This is also suggested by comparing levels of *DMD* transcript as measured by RT-qPCR, using primers up- and downstream of the targeted exons (Supp. Figure 5). As an example, we observe that in control cells, the more efficient AONs skipping exon 51 (Figure 3A), such as PMO-027, −147 and −281 show reduced transcript levels when measuring Exon55-56 (Supp. Figure 5A) in these samples, indicating that there is a correlation between AON efficacy and loss of transcript. This effect was not observed for exon 53 (Supp. Figure 5B).

**Figure 3:**
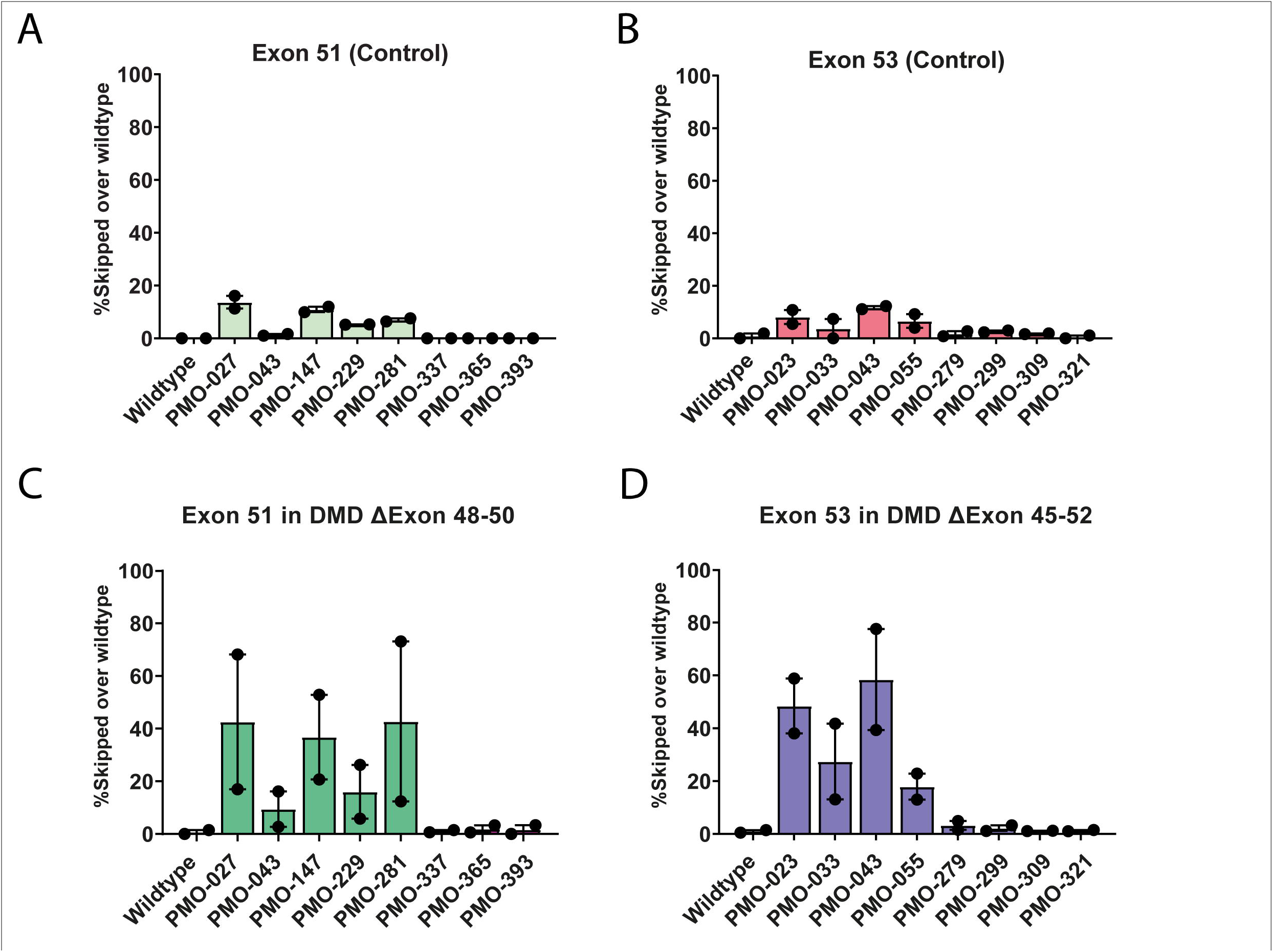
Disruption or restoration of the reading frame due to exon skipping does not change relative compared efficiency. **A**: Exon skipping efficiency of exon 51 of the *DMD* gene in HC myotubes after treatment with exon 51 targeting AONs. Skipping of exon 51 will lead to an out of frame transcript. **B**: Exon skipping efficiency of exon 53 of the *DMD* gene in HC myotubes after treatment with exon 53 targeting AONs. Skipping of exon 53 will lead to an out of frame transcript. **C**: Exon skipping efficiency of exon 51 of the *DMD* gene in DMD patient (ΔExon 48-50) myotubes after treatment with exon 51 targeting AONs. Skipping of exon 51 will lead to an in frame transcript. **D**: Exon skipping efficiency of exon 53 of the *DMD* gene in DMD patient (ΔExon 45-52) myotubes after treatment with exon 53 targeting AONs. Skipping of exon 53 will lead to an in frame transcript.

### Nonsense mediated decay may affect detectable skipping efficiency

We noticed that skipping of exon 51 and exon 53 was more efficient in DMD patient cells, where exon skipping restores the reading frame, than in HC cells, where exon skipping disrupts the reading frame. To test whether NMD might play a role in this observation, both HC and DMD myocytes were nucleofected with PMOs, after which cells were treated with cycloheximide (CHX), an inhibitor of NMD. As a proof of principle, we also included samples nucleofected with an equimolar mix of all 12 available exon 65 targeting PMOs, at the same end concentration (Supp. Figure 6A). The mix of a dozen PMOs leads to minor levels of skipped exon 65, while none of the individual PMOs resulted in detectable levels of exon skipping. Treatment of nucleofected HC cells with CHX causes an increase in detected skipped *DMD* transcript, indicating that NMD might degrade the skipped product, leading to underestimation of skipping efficiency of out-of-frame exons. However, this effect was less pronounced in the DMD exon Δ45-52 cells, and completely absent in the DMD exon Δ48-50 line. RT-qPCR of the *DMD* gene in these samples show a similar, stabilizing effect on the *DMD* transcript, with a modest increase in the total detected expression level (Supp. Figure 6B). Similarly, exon skipping of exon 51 and 53 in the HC (Supp. Figure 6C and 6D) and the two DMD lines (Supp. Figure 6E and 6F) and treatment with CHX yielded inconclusive results. While HC cells showed a slight increase of detectable skipped *DMD* after CHX, DMD cells showed lower detectable skipping efficiency for some PMOs. This result is not dissimilar to previous reports claiming that blocking NMD in DMD cells does not stabilize the *DMD* transcript (García-Rodríguez et al. 2020).

## Discussion

In this study we aimed to elucidate the connection between the efficiency of AON mediated exon skipping, and the splicing characteristics of the introns neighboring the target exon. Our study focused on the *DMD* gene transcripts in human muscle cells, as there is great potential for therapeutic use of AONs in treating Duchenne muscular dystrophy. For this purpose we used a library of 153 custom designed and individually tested *DMD* targeting PMOs with similar physical properties. We found that targeting an exon of which the upstream intron is retained in the transcript longer (a ‘slow intron’) leads to generally higher efficiency of exon skipping than when the exon is preceded by a rapidly spliced (‘fast’) intron. There seems to be no correlation between the splicing speed (‘slow’ nor ‘fast’) of the downstream intron on an exons skippability. Furthermore, we analyzed whether targeting an AON closer to a 5’ or 3’ of either a ‘fast-slow’ or ‘slow-fast’ exon would lead to a difference in exon skipping efficiency. We observed a clear indication that targeting an exon closer to the 5’ end leads to more efficient exon skipping than targeting the 3’ of the same exon. Our study is the first to systematically assess optimal exons and optimal target sites within exons for AON-mediated exon skipping. Other (retrospective) studies on exon skipping efficiency suffered from vastly different physical properties and/or chemistry of the AONs, making comparisons of targeting site efficiency hard to interpret. Our study specifically aimed at reducing differences that could be caused by physical differences in the AONs, by designing PMO AONs of highly similar characteristics such as length, melting temperature, GC content and free energy potential.

Currently, four different *DMD* targeting AONs have been FDA approved for the treatment of Duchenne: Eteplirsen (exon 51), Golodirsen (exon 53), Viltolarsen (exon 53) and Casimersen (exon 45)(Schneider and Aartsma-Rus 2021). Eteplirsen (30-mer) targets nucleotides 66 through 95 of the 233 bp exon 51. Golodirsen (25-mer) and Viltolarsen (21-mer) respectively target nucleotides 36 through 60 or 56 of the 212 bp exon 53. Casimersen’s (22-mer) targeting sequence extends from the last 3 nucleotides of intron 44 through the 19^th^ nucleotide of the 176bp exon 45. Hence, the targeting sites of these therapeutic AONs largely are in accordance with our findings that targeting the 5’ of an exon would be a preferential targeting site over a 3’ targeting AON. However, our AON design never included intronic sequences, so the possibility remains that AONs spanning the intron-exon junction are more efficient than pure exonic targeting AONs.

Our initial screening method only sought to include out-of-frame exons to test our hypothesis on the influence of intronic retention times. While this approach leads to more potential to use the dataset in the development of AONs to restore the *DMD* reading frame in currently untreatable cases, it does lead to the generation of out-of-frame transcripts in the healthy control cell line used. It is therefore likely that nonsense mediated decay could degrade skipped transcripts, leading to an underestimation of exon skipping potential of an NMD-triggering AON. Indeed, we observe that inhibition of NMD by CHX treatment in HC cells treated with exon 51 or 53 targeting PMOs occasionally increases the observed amount of exon skipping as determined by RT-PCR. Intriguingly, the *DMD* stabilizing effect was not observed in muscle cells from DMD patients that would be amenable to exon skipping therapies with exon 51 or 53 targeting PMOs. In these cells, while skipping levels were markedly higher than in HC cells, no additional benefit was seen from a combination of CHX and AONs, suggesting that the *DMD* transcript in DMD patients is not actively degraded through NMD as was reported previously(García-Rodríguez et al. 2020).

The exons included in this study were positioned over the entirety of the *DMD* gene as far as our described design constraints allowed so. Transcription of *DMD* is suggested to take as long as 16 hours, and there is a reported imbalance between the abundance of the 5’ and 3’ end of the transcript(Spitali et al. 2013). The 3’ of *DMD* being underrepresented indicates that potentially, transcription is properly initiated but unfinished. As splicing of *DMD* occurs co-transcriptionally, there might be a correlation between the transcription of the RNA molecule and the possibility to skip an exon, as a proximal exon would be more abundantly present in cells. However, we did not see a correlation between exon skipping efficiency of our tested PMOs and the location of the exon in the gene. E.g.: the skippability of exon 17 and 70 is relatively high, while the skippability of exon 10 and 65 was low.

The splicing maps indicating the splicing order of introns of the *DMD* gene were based on results we generated in HC cells (Gazzoli et al. 2016). It is yet unclear what properties make an intron ‘slow’ or ‘fast’. If this information is contained within the intron itself, it is possible that a large deletion encompassing part of an intronic region will alter the splicing characteristic of the remaining flanking exons. That is, an exon originally flanked by a slow intron might be flanked by a fast intron upon genomic rearrangement in a DMD patient. Whether this occurs would most likely highly depend on the exact breakpoint of the mutation, which is often unique in each patient (Aartsma-Rus et al. 2006; Bladen et al. 2015). Whether this ‘intronic-splice-speed-switching’ actually occurs in DMD and whether this would have consequences for AON mediated therapeutics remains unknown until the nature of ‘slow and ‘fast’ introns are elucidated further. DMD patient cells with the same deleted exon, but carrying unique intronic rearrangements, could be treated with an AON library as we present here to determine exon skipping efficiency. Overlaying the intronic deletion pattern with exon skippability might then lead to the discovery of new motifs in the intron, which could be highly valuable information in clinical and pre-clinical research on Duchenne therapies, and fundamental understanding of the splicing process.

In conclusion, our data provide new insights into preferred selection of exon skipping target exons, which can be applied when selecting the next exon skipping therapy candidate in DMD and other applicable disease. Moreover, we show in an unbiased manner that generally, the optimal exon targeting region is the 5’, which can guide future design for AON based exon skipping therapies.

## Material and methods

### Cell culture

Healthy control (HC (KM155)) and DMD patient cells (8036 and 6311) were kindly provided by Prof. Vincent Mouly (Institute de Myology) and have been described before (Mamchaoui et al. 2011; Echigoya et al. 2017; Echigoya et al. 2019). Cells were maintained in skeletal muscle growth medium (Promocell C-23060), supplemented with 15% heat-inactivated fetal bovine serum (HI-FBS) and 50 μg/ml gentamycin. For differentiation of proliferating myoblasts to myotubes, medium was replaced with fusion media consisting of Dulbecco’s modified eagle medium (DMEM)-Glutatamax, high-glucose (Gibco 61965059), supplemented with 2% HI-FBS and 1% penicillin-streptomycin (Gibco 15140-122). Cells were maintained in humidified incubators at 37°C and 5% CO_2_. Cells were regularly tested for mycoplasma infection by use of the mycoalert (Lonza, LT07-318) kit.

### AON design

PMOs were designed to specifically target selected *DMD* exons, while exhibiting similar physical properties such as G/C content, number of nucleotides and melting temperature. For determination of the free energy of potential secondary structures (intramolecular and homodimer formation), RNAstructure 6.2 was used (Reuter and Mathews 2010). Each potential 24/25-mer AON targeting the 47 *DMD* exons outlined below was designed and analysed (∼11.213 AONs in total). The average G/C content and melt temperature were calculated for the entire set, and AONs that deviated >1x standard deviation were excluded. Exons where less than 25 potential AONs remained were excluded from further consideration, as diversity of targeting sites would be too low. PMOs were synthesized and quality controlled in-house at Sarepta Therapeutics inc. Cambridge, MA, USA. PMOs were dissolved in sterile saline and diluted as stocks of 1 mM. Sequences and properties of all PMOs are listed in supplementary table 1.

### Electroporation

For delivery of PMOs to myoblasts cells, a Lonza Amaxa 4D nucleofector with accompanying X unit was utilized, as described in (Goossens and Aartsma-Rus 2022). In brief, cells were trypsinized and resuspended in nucleofection P1 buffer at a concentration of 1*10^6^ cells per 20 μl of buffer. Then, 20 μl of P1 cell suspension was transferred to each well of the 16 well nucleocuvette, after which 1 μl of 1 mM PMO was added to the well, for a final concentration of 50 μM. Cells were electroporated in the X unit using program CM-137, and allowed to recover for 10 min at room temperature (RT). After recovery, cells were carefully resuspended in 200 μl growth medium and transferred to 6-well plates containing equilibrated growth medium. Cells were grown for at least 48h before confluent cultures were allowed to differentiate to myotubes for 72h as described.

### RNA isolation and cDNA synthesis

For RNA isolation, myotubes in each well were lysed in 500 μl Tri-sure lysis reagent, after which 200 μl chloroform was added and phases were separated by centrifugation at 16.200 relative centrifugal force (RCF) for 15 min at 4°C. The aqueous phase was transferred to 500 μl of 2-propanol and RNA was precipitated by centrifugation for 15 min at 4°C at 16.200 RCF. The pellet was washed twice with 70% ethanol, air-dried and resuspended in 25 μl RNAse-free milli-Q (MQ). RNA concentration and purity was determined using a ND-1000 Nanodrop (thermo-scientific) and provided as A_260_/A_230_ and A_260_/A_280_ ratios. For cDNA synthesis, 1 μg of total RNA was used using bioscript Tetro (bioline BIO-65050), according to the manufacturer’s instructions in a 20 μl reaction. Samples were diluted to 100 μl final volume with MQ.

### RT-PCR exon skipping analysis

RT-PCR analysis was used to determine *DMD* exon skipping efficiency. Ten percent of generated cDNA was used per reaction using specific sets of intron spanning primers (See supplementary table 2), and Dreamtaq polymerase (Thermo scientific EP0713). Reactions of 25 μl total consisted of 2.5 μl 10x reaction buffer (green), 0.2 μl DreamTaq polymerase (1 Unit), 1 μl 10 μM forward primer, 1 μl 10 μM reverse primer, 1 μl DNTP mix (10 μM each nucleotide) and 9.3 μl MQ. Amplification was performed in T-100 thermal cyclers (Bio-Rad) using the following parameters: 1: 95°C 2 min,; 2: 95°C 30 sec.; 3: 60°C 30 sec.; 4: 72°C 45 sec.; 5: Go to step 2, 34 additional cycles; 6: 72°C 5 min. A fraction of each sample was analyzed using standard agarose TRIS-Borate-EDTA (TBE) gel-electrophoresis. DNA quantity in the PCR samples was assessed using Qubit dsDNA broad range reagent, measured in a Spectramax ID3 instrument. Samples were diluted to 0.2 ng/μl and analyzed using an Agilent Femto Pulse with NGS separation gel. Peaks were called using the accompanying Prosize Data analysis software version 4.0.2.7. Skipping efficiency was calculated by determining the ratio of concentration (in nmole/L) of the skipped product over the total concentration (in nmole/L) of the sum of the full-length and skipped product. Statistical analysis was performed in Graphpad Prism Version 8.

### RT-qPCR analysis

For gene expression analysis, we used 2% of the generated cDNA samples described above per reaction. The RT-qPCR reaction consisted of 4 μl SensiMix 2x SYBR master mix (bioline QT605-05), 2 μl cDNA, 1 μl forward primer (10 μM) and 1 μl reverse primer (10 μM) (see supplementary table 2). Each sample/primer combination was measured in a technical triplicate. Samples were manually pipetted in 384-well plates (Framestar 480/384, 4ti-0381 4titude) and run in a CFX-384 Real-time PCR system (Bio-Rad). Cycling conditions were as follows: 1: 95°C 5:00; 2: 95°C 0:10; 3: 60°C 0:30 (plate read); 4: Go to step 2 39 additional times; 5: melt curve 60°C to 95°C, 0.5°C increase per cycle of 0:05. Data was analyzed using the CFX-Maestro software version 2.0, with baseline and Cq value calculation set to ‘auto calculation mode’. CFX-Maestro was also used to determine run quality by studying the melt curves generated for each product, as well as internal QC functions. Expression values were normalized to the housekeeping genes *GAPDH* and *GUSB*, using the ΔΔCt method, after which statistical analysis and visualization was performed using Graphpad Prism Version 8.

## Acknowledgements

We thank Dr. Vincent Mouly (Institute of Myology, Paris, France) for providing the myogenic cell lines used in the study. Authors are part of the European BIND consortium. Several authors of this publication are members of the Netherlands Neuromuscular Center, the Duchenne Centrum Netherlands (funded by Spieren voor Spieren) and the European Reference Network for rare neuromuscular diseases EURO-NMD.

## Supp. Figures

**Supp. Figure 1:**

Validation of PMO delivery to immortalized HC myoblasts by electroporation

**A**: Relative expression of various genes measured by RT-qPCR after nucleofection of an exon 51 targeting PMO with Lonza buffer systems and Amaxa pulse programs as indicated. The *DMD* exon 50-52F_52R primer set is specific to measure the presence of the *DMD* gene when exon 51 is skipped, while the exon 49-50 primer set shows the presence of both skipped and unskipped *DMD. MYOG* and *MYH3* are measured as an indication for myogenic proliferation. Data is normalized to housekeeping genes *GUSB* and *GAPDH*.

**B**: Comparison of the same set of RT-PCR samples measuring exon 51 skipping percentages as the ratio of skipped over total (skip+full-length) PCR product, derived from a set of HC myotubes after nucleofection with an exon 51 targeting PMO. Adjacent bars show the same PCR product measured on either system as indicated.

**Supp. Figure 2:**

Separated skipping data presented in figure 1. Error bars indicate SD of two independent samples.

**Supp. Figure 3:**

**A**: Separated skipping data presented in figure 2B. Error bars indicate SD of two independent samples.

**B**: Separated skipping data presented in figure 2C. Error bars indicate SD of two independent samples.

**C**: Data presented in figures 2B and 2C was reanalyzed and grouped to separate exons based on their upstream and downstream intron class as indicated.

**D**: Data presented in figures 2B and 2C was reanalyzed and grouped to separate exons based on whether efficient skipping of the AON would lead to an in frame or out of frame transcript as indicated.

**Supp. Figure 4:**

Separated skipping data presented in figure 2. Error bars indicate SD of two independent samples.

**Supp. Figure 5:**

RT-qPCR expression of DMD levels in samples presented in figure 3. Measurements for the *DMD* transcript using a primer set upstream (Ex38-39) and downstream (Ex55-56) of the skipped exons are shown. *MYH3* expression is used as an indication of myogenic proliferation in the sample. Data is normalized to housekeeping genes *GUSB* and *GAPDH*.

**Supp. Figure 6:**

**A**: Exon skipping efficiency of HC and DMD cells nucleofected with no PMOs (WT) or a mix of 12 exon 65 targeting PMOs, and treated with DMSO or CHX as indicated.

**B**: RT-qPCR expression of DMD levels in samples presented in figure 6A. Measurements for the *DMD* transcript spanning exon 38-39 and exon 55-56 are shown. *MYH3* expression is used as an indication of myogenic proliferation in the sample. Data is normalized to housekeeping genes *GUSB* and *GAPDH*.

**C**: Exon skipping efficiency of exon 51 of the *DMD* gene in HC myotubes after treatment with exon 51 targeting AONs and CHX as indicated.

**D**: Exon skipping efficiency of exon 53 of the *DMD* gene in HC myotubes after treatment with exon 53 targeting AONs and CHX as indicated.

**E**: Exon skipping efficiency of exon 51 of the *DMD* gene in DMD patient (ΔExon 48-50) myotubes after treatment with exon 51 targeting AONs and CHX as indicated.

**F**: Exon skipping efficiency of exon 53 of the *DMD* gene in DMD patient (ΔExon 45-52) myotubes after treatment with exon 53 targeting AONs and CHX as indicated.

